# A conceptual view at microtubule plus end dynamics in neuronal axons

**DOI:** 10.1101/062711

**Authors:** André Voelzmann, Ines Hahn, Simon P. Pearce, Natalia Sánchez-Soriano, Andreas Prokop

## Abstract

Axons are the cable-like protrusions of neurons which wire up the nervous system. Polar bundles of microtubules (MTs) constitute their structural backbones and are highways for life-sustaining transport between proximal cell bodies and distal synapses. Any morphogenetic changes of axons during development, plastic rearrangement, regeneration or degeneration depend on dynamic changes of these MT bundles. A key mechanism for implementing such changes is the coordinated polymerisation and depolymerisation at the plus ends of MTs within these bundles. To gain an understanding of how such regulation can be achieved at the cellular level, we provide here an integrated overview of the extensive knowledge we have about the molecular mechanisms regulating MT de/polymerisation. We first summarise insights gained from work *in vitro*, then describe the machinery which supplies the essential tubulin building blocks, the protein complexes associating with MT plus ends, and MT shaft-based mechanisms that influence plus end dynamics. We briefly summarise the contribution of MT plus end dynamics to important cellular functions in axons, and conclude by discussing the challenges and potential strategies of integrating the existing molecular knowledge into conceptual understanding at the level of axons.

## 1. Introduction

Axons are the slender, up-to-a-metre long processes of neurons which form the cables that wire the nervous system. These delicate structures often need to survive for a lifetime, i.e. many decades in humans. Parallel polar, predominantly plus end out oriented, bundles of microtubules (MTs) run all along axons to form their structural backbones and highways for life-sustaining transport (Baas and Lin, 2011). These MT bundles are so densely packed, with MTs being separated by less than 100nm, that they can only be reliably resolved through electron microscopy (Mikhaylova et al., 2015).

MTs are important at every life stage and for virtually all morphogenetic changes of axons (Penazzi et al., 2016; Prokop, 2013) (Fig. 1). Alterations in MT properties are often linked to developmental and/or intellectual brain disorders, and the precocious decay of MT bundles is seen as an important cause for axon degeneration (Adalbert and Coleman, 2012; Neumann and Hilliard, 2014). MTs are therefore viewed as attractive targets for drug therapies (Baas and Ahmad, 2013; Benitez-King et al., 2004; Eira et al., 2016).

**Figure 1:**
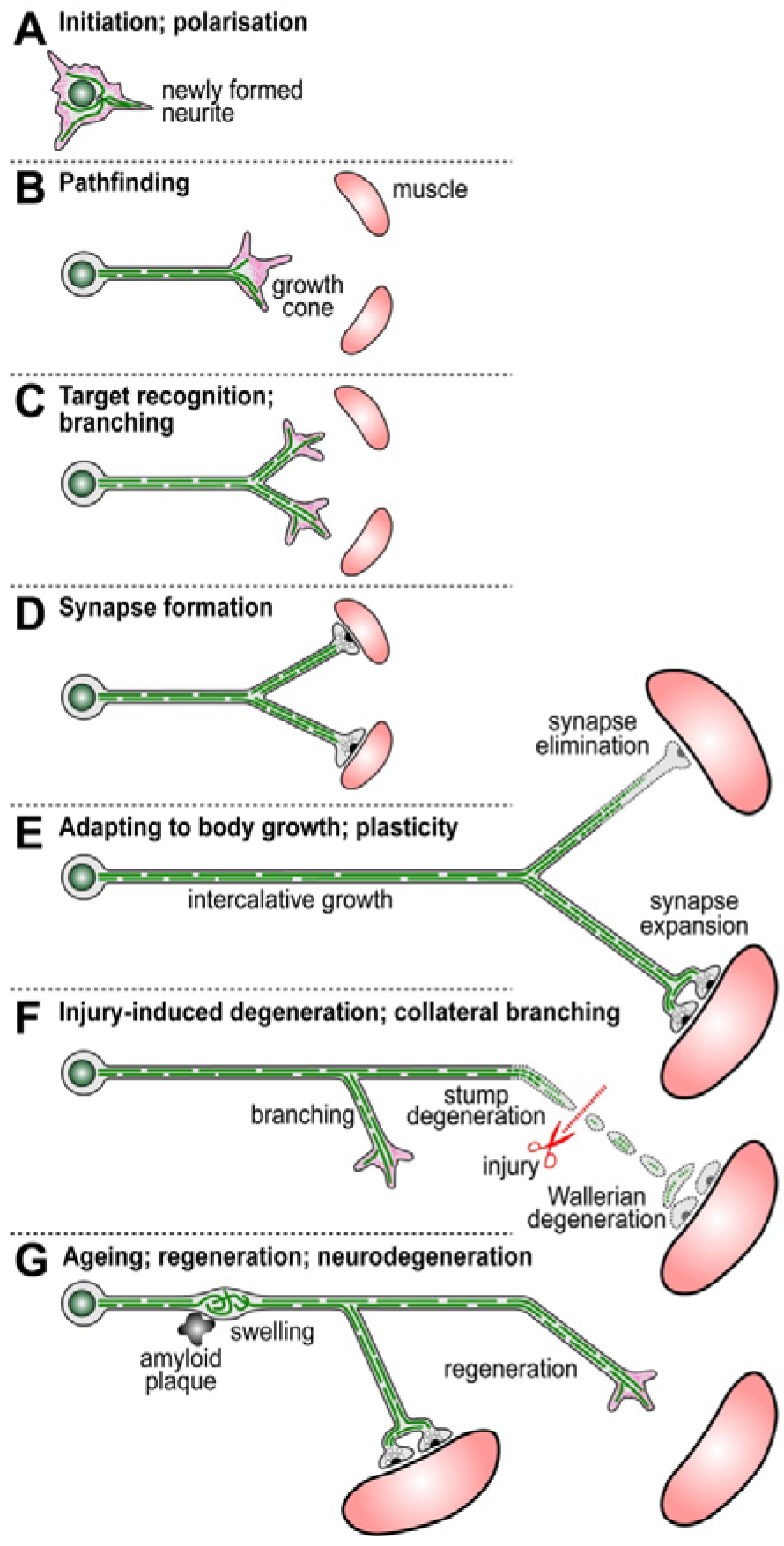
Life stages of axons. Images show one neuron during its various life stages, with the cell body on the left and axon on the right; MTs are shown as green interrupted lines, actin in magenta and muscles as pink round shapes. The different life stages are indicated in the headings, and the various biological processes associated with those stages are annotated in each figure.

*In vitro* and *in vivo*, MTs are dynamic, characterised by cycles of endothermic polymerisation, and exothermic depolymerisation (Brouhard, 2015) (section 3). However, *in vivo*, these dynamics are not left to chance but are controlled by different classes of MT binding proteins (MTBPs) and the proteins they recruit and associate with. Such MT binding and associating proteins regulate polymerisation, severing, depolymerisation, stability, transport and force production, as well as cross-linkage and interaction with other organelles or cell structures (Penazzi et al., 2016; Prokop et al., 2013).

Many of these MTBPs and associated proteins have acknowledged links to brain disorders or other human diseases (Prokop et al., 2013). Although the molecular function of these proteins is often known, this knowledge is usually not sufficient to explain their associated diseases. One key challenge is posed by the fact that MT regulators form complex networks that establish robust and redundant machinery which often responds with surprisingly subtle changes upon mutation of single components. Furthermore, the cytoskeleton is required for virtually every cell function including cell division, dynamics and shape, adhesion and signalling, intracellular trafficking, and organelle biology (Alberts et al., 2014). Therefore, mutations of cytoskeletal regulator genes can be expected to have pleiotropic effects, further complicating their study and requiring a broader spectrum of functional readouts.

To shed light into these complex networks and to find explanations for disease phenotypes, it seems a promising strategy to dissect cytoskeletal machinery into its conceptual sub-machineries and then study how these sub-machineries interface with each other. Along these lines, a previous review was dedicated to the intricate relationship of motor proteins and MTs (Prokop, 2013). Here we provide a conceptual overview of the sub-machinery (or sub-machineries) that governs the polymerisation and depolymerisation of MTs in axons.

## 2. The importance of MT dynamics in axons

In axons, MTs primarily grow in the anterograde direction with a certain bias towards the more distal axon segment, but retrograde polymerisation events have also been reported when performing live imaging with typical plus end markers (Kollins et al., 2009; Sánchez-Soriano et al., 2010; Stepanova et al., 2003; Yau et al., 2016). These may reflect either plus end polymerisation of antiparallel MTs or minus end polymerisation of parallel MTs, which can be observed *in vitro* (Akhmanova and Hoogenraad, 2015; Akhmanova and Steinmetz, 2015; Jiang et al., 2014).

The abundance of MT polymerisation along the axon shaft could mean that axonal MT mass is generated in the shaft, rather than in growth cones (Fig.1C) where polymerisation gives rise to rather short-lived MTs similar to those in non-neuronal cells (Prokop, 2013; Prokop et al., 2013). It has been proposed from studies in developing vertebrate and *Drosophila* neurons that the MT mass generated in the axonal shaft gradually shifts anterogradely (Miller and Sheetz, 2006; Roossien et al., 2013). In such a scenario, the growth potential would originate from MT bundles along the axon and growth cones would be the intelligent units which can control this extension and its directionality.

This model of shaft-based growth is further supported by the fact that significant axon growth occurs at post growth cone stages (Fig. 1E). Thus, many growth cones reach their target destination and transform into synapses already in the embryo. Since the organism continues to grow towards its much larger adult dimension, significant further axon elongation has to occur through intercalative shaft growth (Bray, 1984).

Even in adulthood, MT polymerisation does not seize. Firstly, it is required for plastic growth and branching as well as for regenerative growth (Fig. 1F, G) (Bilimoria and Bonni, 2013; Bradke et al., 2012; Kalil and Dent, 2014; Lewis et al., 2013). Secondly, most axons have to be sustained for the lifetime of an organism, i.e. up to a century in humans, and continued MT turnover is needed to prevent senescence through mechanical wear (Dumont et al., 2015) and remove irreversible posttranslational modifications (Janke and Kneussel, 2010). Accordingly, continued MT polymerisation has been demonstrated in adult *Drosophila* brains (Medioni et al., 2015; Nguyen et al., 2011; Rolls et al., 2007; Soares et al., 2014) and similar results can be expected in vertebrates. Thirdly, the MT volume of many axons seems to be a well-regulated property. This is suggested by neuron type-specific axon diameters and MT densities (Wortman et al., 2014), and the fact that artificially slimmed axons (achieved through pulling) reconstitute their original diameters (Bray, 1984). How axon diameters are sensed and MT volume (i.e. number and density) maintained within, remains largely unresolved. Intermediate filaments may play a role in regulating axon diameter and MT volume (Kriz et al., 2000) (and references therein). However, other mechanisms must be in place, because *Drosophila* also has defined axon diameters but lacks intermediate filament genes (Weber et al., 1991).

In any case, it seems a reasonable assumption to propose that axons “know” and continuously adapt their MT volume, and that the MT de/polymerisation machinery has to be one key element of this regulation. This poses the fundamental question of what the molecular nature of this machinery is.

## 3. Properties of MTs and MT dynamics as revealed by *in vitro* studies

Before delving into the molecular complexity observed at the cellular level, it is important to provide a brief overview of our knowledge about the physical and biochemical properties of tubulin and MT dynamics gained from 5 decades of intense *in vitro* studies.

MTs are the stiffest of the three cytoskeletal polymers with a persistence length of ∼5 mm, as compared to ∼12 μm measured for actin filament (Fletcher and Mullins, 2010; Howard, 2001). The building blocks of MTs are α- and ß-tubulin heterodimers which will polymerise into MTs even *in vitro*. They form strong longitudinal head-to-tail bonds which make up the protofilaments. As detailed elsewhere (Chretien and Fuller, 2000; Howard, 2001), between 8 and 19 protofilaments arrange in parallel into a sheet by forming lateral bonds between homologous (α-α, ß-ß; B-type lattice) or heterologous subunits (α-ß; A-type). To form a tube, the edges of this sheet join, forming a seam at which the joining protofilaments are out of register by 2-4 tubulins (referred to as “start helical arrangement”; Fig. 2). In consequence, the lateral bonds of MTs take on a helical structure where the angle of the helical rise (as a function of start helical arrangement and protofilament number) can cause a skew of lateral bonds of up to 2°; such skew angles were proposed to be energetically unfavourable (Chretien and Fuller, 2000). Accordingly, the majority of MTs contain 13-protofilaments with a 3-start helical arrangement, where the skew angle is almost 0° and the outer diameter is 25nm. However, MTs with different protofilament numbers occur *in vivo*, but the functional relevance of this is not clear (Chrétien and Wade, 1991) (and references therein).

**Figure 2:**
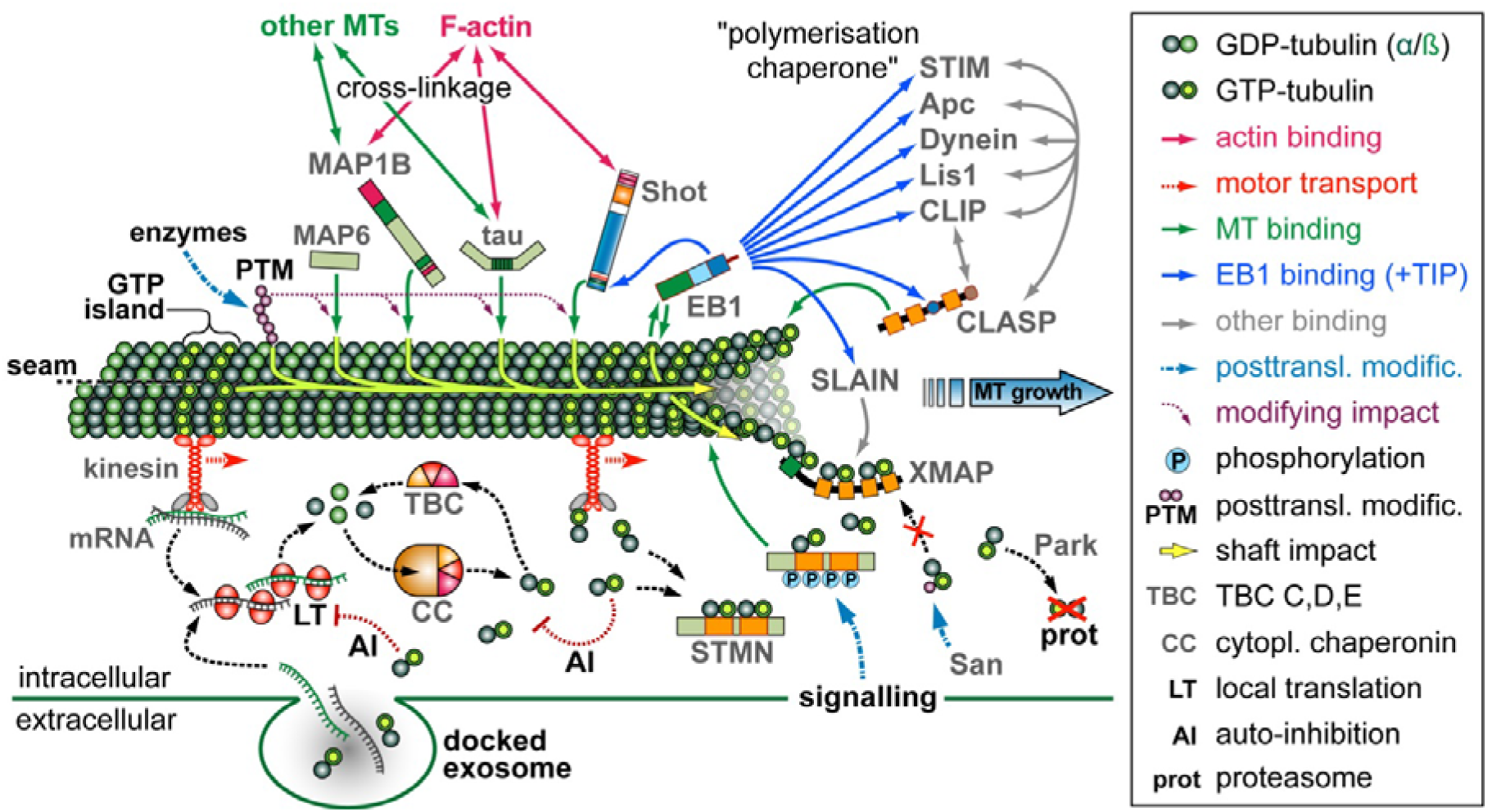
Integrated view of the various mechanisms regulating MT plus end dynamics. The image shows a single MT surrounded by regulating factors (gene/protein names in grey). Symbols are explained in the box on the right and further aspects are annotated in the figure, with details provided in the main text.

MTs are polar, with a plus end (with exposed ß-tubulin) and a minus end (with exposed α-tubulin). *In vitro*, both ends have very different kinetics of MT polymerisation or depolymerisation; as an extreme example, conditions can be set so that MTs undergo treadmilling similar to actin filaments, polymerising at plus ends and depolymerising at minus ends (Margolis and Wilson, 1998; Panda et al., 1999).

*In vitro*, new MTs are either nucleated via stabilised MT fragments anchored to the glass surface as powerful ‘seeds’ to form MTs (Akhmanova and Steinmetz, 2015) or polymerisation is facilitated by additional factors such as the MTBP tau (Akhmanova and Steinmetz, 2015; Brandt and Lee, 1993a, b) This resembles the situation in cells, where new MTs originate from nucleation complexes; these complexes contain a ring of □-tubulin as the key component onto which tubulin heterodimers are polymerised, and which subsequently anchors the minus ends of the newly formed, elongating MT. Nucleation complexes containing □-tubulin can be found on centrosomes, the kinetochore of chromosomes, other MTs, the nuclear envelope, the Golgi surface or the cell cortex (Petry and Vale, 2015; Teixido-Travesa et al., 2012). Accordingly, □-tubulin is also found in axons and, together with the augmin complex, forms a plus end-out directed nucleation machinery that can uphold MT bundle polarity (Nguyen et al., 2014; Sanchez-Huertas et al., 2016; Stiess et al., 2010). As a further mechanism, MT fragments were suggested to act as sites for re-polymerisation in axons (Baas et al., 2016). Such MT fragments could potentially be bound and directionally anchored to the axonal cortex via gigantic actin-MT linker molecules called spectraplakins (Baas et al., 2016; Nashchekin et al., 2016) (section 5).

Subsequent polymerisation *in vitro* can also be achieved in the absence of MTBPs, but polymerisation speeds are significantly lower than observed *in vivo* or when adding the right cocktail of MTBPs (Zanic et al., 2013). In MTBP-free *in vitro* assays, the essential parameters determining the polymerisation speed include the concentrations of tubulin, guanosine triphosphate (GTP), Mg^2+^(influencing the GTP-tubulin interaction), calcium chelators, as well as an appropriate temperature to drive the endothermic polymerisation reaction (usually 37°C); also the tubulin isotype has an impact, with β3-tubulin de/polymerising considerably slower than other β-tubulins (Grover and Hamel, 1994; Lu and Luduena, 1994b).

The addition of GTP is important because α- and β-tubulins bind GTP (referred to as GTP-tubulin). Under standard conditions, GTP-tubulin is effectively incorporated into MTs, whereas GDP-tubulin incorporation *in vitro* requires non-physiological conditions (Lin and Hamel, 1987) (and references therein). Following incorporation into MTs, the GTP of β-tubulin (but not α-tubulin) gets hydrolysed (these heterodimers are referred to as GDP-tubulin) (Akhmanova and Steinmetz, 2015). This hydrolysis occurs with a certain delay, so that GTP-tubulin forms a zone of several hundred nm from the polymerising MT tip which is often referred to as the GTP cap (Fig. 2). As detailed elsewhere (Akhmanova and Steinmetz, 2015), *in vitro* experiments have demonstrated that the GTP cap has two important functions. Firstly, it provides the substrate to which EB proteins bind which, in turn, recruit a huge number of further proteins to the MT plus end (see section 4 for details). Secondly, the GTP cap directly stabilises MT plus ends as a consequence of its structural configuration which is favourable for the alignment into straight protofilaments. However, the hydrolysis to GDP-tubulin introduces a compaction within the tubulin molecule which seems to generate longitudinal strain and hence stored elastic energy (Brouhard and Rice, 2014). This has no immediate consequence, for as long as lateral and longitudinal bonds hold the tubulin in its straight confirmation. But if the GTP cap shrinks, GDP-tubulin protofilaments may break their lateral bonds and release their stored energy by peeling off like banana skin, thus very rapidly depolymerising the MT plus end. These dynamic changes at MT plus ends (termed dynamic instability) occur *in vitro* and in cells, and are referred to as pause (de/polymerisation comes to a halt), catastrophe (polymerisation/pause turns into depolymerisation) and rescue (depolymerisation turns into pause or repolymerisation).

Finally, MTs *in vitro* do not only de/polymerise at their plus ends, but also at their minus ends (though with far slower kinetics) (Akhmanova and Hoogenraad, 2015; Hendershott and Vale, 2014), and it has been shown that polymerisation can occur even along the MT lattice to repair lateral damage (Schaedel et al., 2015).

## 4. Regulating tubulin availability in axons

As has become clear from *in vitro* studies, the concentrations of tubulin dimers at plus and minus ends of MTs are essential parameters determining de/polymerisation (section 3). This raises the important question of how tubulin pools are controlled in axons.

In neurons, tubulin is under tight transcriptional control (Blackmore, 2012) and synthesis of specific isotypes can be favoured by stabilisation of their mRNAs (Rogers et al., 1993). Once produced, tubulin proteins tend to be rather stable, with half-lives ranging between 15-50 hrs depending on the cellular model (Caron et al., 1985a; Mooney et al., 1994; Semotok and Lipshitz, 2007). However, this half-life is not absolute but subject to tissue type (Burow et al., 2015), culture substrate (Mooney et al., 1994) and cell-cycle state (Ducommun et al., 1990). Furthermore, tubulins can be recycled for new rounds of polymerisation (Nogales and Wang, 2006), as illustrated also by *in vitro* studies with purified brain tubulin which can be repeatedly polymerised and depolymerised (e.g.) (Chretien and Fuller, 2000).

How the pools of tubulin are supplied and regulated in axons, especially during times of high net demand (e.g. during axon growth), remains little understood. One mode is motor protein-mediated transport of tubulin mRNAs, monomers, dimers or even MT fragments travelling at fairly low speeds of ∼1mm per day (Fig. 2) (Frühbeis et al., 2013; Galbraith and Gallant, 2000; Tang et al., 2013) For a more immediate supply, axons contain the machinery to perform local tubulin biosynthesis from mRNAs, although cell culture studies have suggested that not more than 5% of axonal tubulin are generated this way (Eng et al., 1999; Giuditta et al., 2002b; Giuditta et al., 2008; Jung et al., 2012; Lee and Hollenbeck, 2003). However, it might be expected that local provision is far more essential in long axons *in vivo*, where also glia-derived supply of tubulin mRNA and/or protein, for example via exosomes, may play important roles (Fig. 2) (Frühbeis et al., 2013; Giuditta et al., 2002a). As a further advantage, local biogenesis would provide flexible means to fine-tune tubulin levels, and posttranscriptional cytoplasmic auto-regulation of tubulin has been reported (Theodorakis and Cleveland, 1992) (and references therein). For example, artificially raising the levels of tubulin dimers in de-nucleated fibroblasts (a situation similar to axons) caused a down-regulation of tubulin biosynthesis, reducing both mRNA and protein half-life (Fig. 2) (Caron et al., 1985b).

Tubulin levels in neurons can also be regulated through the ubiquitin ligase parkin and proteasome-mediated degradation (Fig. 2) (Ren et al., 2003), and through components of the chaperone machinery (Fig. 2). The biogenesis of GTP-tubulin is a multi-step process where native tubulins first encounter prefoldin, then enter the large ATP-dependent cytoplasmic chaperonin complex where they acquire their shape and GTP-bound status, after which the chaperonin cofactors A-E (TBCA-E) take over to catalyse dimerisation and cytoplasmic release (Bhamidipati et al., 2000; Tian et al., 1996). However, TBCC, -D and -E can also act outside this machinery by binding and cleaving heterodimers (Fig. 2) (Bhamidipati et al., 2000; Serna et al., 2015). This fine balance between MT promoting and destabilising roles of these TBCs may explain the contradictory results obtained from different gain- and loss-of-function experiments with TBCs, suggesting them to be either promoters or inhibitors (Bhamidipati et al., 2000; Bömmel et al., 2002; Jin et al., 2009) (and references therein).

It appears therefore that free cytoplasmic tubulin heterodimers would not have a long half-life and require stabilising factors. Such stabilisers could be stathmins which contain two tubulin binding regions (four in some isoforms of *Drosophila* stathmin), each binding one tubulin heterodimer (Fig. 2) (Chauvin and Sobel, 2015; Lachkar et al., 2010). Whilst stathmins are traditionally considered sequestering factors for tubulin heterodimers and inhibitors of MT polymerisation, their reported mutant or over-expression phenotypes suggest a complicated mix of MT promoting and inhibiting roles (Chauvin and Sobel, 2015; Manna et al., 2007; Manna et al., 2009; Riederer et al., 1997), as similarly observed for TBCs. These findings are likely due to the opposing roles that stathmins play (Chauvin and Sobel, 2015; Gupta et al., 2013; Nouar et al., 2016) (and references therein). On the one hand, stathmins are likely to maintain pools of tubulin heterodimers, protecting them from degradation and withholding them from autocatalytic down-regulation of their own biosynthesis, as was explained above. On the other hand, they induce catastrophes through active binding to protofilaments at MT plus ends and withdrawing tubulin heterodimers from the polymerisation machinery. These various functions are controlled by four phosphorylation sites in the Stathmin Like Domain (SLD) which are targeted by various signalling pathways (Fig. 2). A key question is therefore how phosphorylation and heterodimer release of stathmins is spatiotemporally controlled in axons, and how this is coordinated with polymerase activity (for example whether there is close functional collaboration as is the case for Enabled and profilin during actin polymerisation) (Bear and Gertler, 2009).

Apart from tubulin availability in the axon, also the type of tubulins needs consideration, because different tubulins have significantly different polymerisation dynamics (section 3), suggesting neuron type-specific deviations. Furthermore, the acetyltransferase Nat13/San was shown to mediate acetylation of cytosolic tubulin heterodimers at K252 of ß-tubulin, thus inhibiting their incorporation into MTs (Fig. 2) (Chu et al., 2011).

Taken together, a cocktail of mechanisms regulates the availability of tubulins, thus influencing MT de/polymerisation in neurons. The importance of this machinery is clearly illustrated by the prominent links that components of this machinery have to brain disorders.

## 5. The MT de/polymerisation machinery at MT plus ends

The number of proteins associating with MT plus ends (referred to as +TIPs) is steadily increasing, and many have been reported to influence MT stability or polymerisation (Fig. 2) (Akhmanova and Steinmetz, 2010; Akhmanova and Steinmetz, 2015; van de Willige et al., 2016). Strikingly, no reports so far seem to have pinpointed a single master +TIP of polymerisation, the absence of which would cause the total or next to complete loss of MTs. As one explanation, the “polymerisation chaperone hypothesis” was put forward (Fig. 2) (Gupta et al., 2014). This hypothesis states that the many known functional interactions between the various +TIPs construct molecular networks that culminate in a protein superstructure protecting and stabilising MT plus ends and promoting their polymerisation. Loss of one or few types of +TIPs from such superstructures would have little impact due to functional redundancy.

Alternatively, there could be two or more central players which can partly compensate for one another. The other +TIPs would have less significant functions such as fine tuning plus end polymerisation dynamics and adapting +TIP dynamics to different cellular contexts (see also section 7). As will be discussed, central players could be EB proteins and XMAP215, and potentially CLASP. In support of this notion, *in vitro* studies demonstrated a strong synergy between EB1 and XMAP215, significantly speeding up MT polymerisation to *in vivo*-like levels (Zanic et al., 2013).

As summarised elsewhere (Akhmanova and Steinmetz, 2015), up to a few hundred EB molecules can bind to a MT plus end with a very short dwell time, i.e. in a highly dynamic manner with constant turn-over. EB localisation at polymerising MT plus ends closely correlates with the GTP cap, mediated by the N-terminal calponin homology domain which simultaneously binds to four GTPs, linking together tubulin heterodimers across two parallel protofilaments (Akhmanova and Hoogenraad, 2015). In this way, EBs might stabilise lateral and longitudinal tubulin bonds. At the same time, EBs also promote GTP hydrolysis so that, at high concentration, they may cause catastrophes - representing a further set of factors juggling a balance between promotion and inhibition at MT plus ends (Akhmanova and Hoogenraad, 2015; Brouhard and Rice, 2014; Maurer et al., 2014).

The importance of EBs for neuronal microtubules was demonstrated in *Drosophila*, where EB depletion causes severe MT disorganisation and severe axon shortening (Alves-Silva et al., 2012). Short axons are likely to reflect a significant reduction in MT volume due to reduced net-polymerisation. This could either be explained through the direct effects of EB proteins on the MT lattice, or through the role of EBs as scaffold proteins recruiting numerous +TIPs. Thus, the C-terminal coiled-coil and EB homology domains of EB proteins mediate homo- and hetero-dimerisation as well as interaction with other +TIPs, allowing those proteins to “hitchhike” on MT plus ends (Fig. 2) (Akhmanova and Steinmetz, 2015).

These MT plus end hitchhikers often have a certain ability to bind MTs directly (a property that, combined with EB binding, could enable them to “surf” along the growing MT plus ends) (Gupta et al., 2014). They also form numerous links with one another, which has led to the before mentioned chaperone hypothesis (Gupta et al., 2014). However, their localisation at MT plus ends is essentially determined by direct interaction with the C-terminus of EB proteins: either via a CAP-Gly domain binding to the EEY tail (e.g. CLIPs and p150), or via positively charged protein stretches containing SxIP motifs which bind to the EB homology domain (e.g. APC, spectraplakins, CLASP, KIF2C, KIF11, NAVs and SLAIN) (Akhmanova and Steinmetz, 2015; Honnappa et al., 2009). The C-terminus of EBs is very short so that binding of SxIP motifs and CAP-Gly domains to EB proteins is considered mutually exclusive and hence competitive (Duellberg et al., 2014). Binding to these specific domains seems to occur in hierarchical order of affinity which can also be modified through signalling events (Akhmanova and Steinmetz, 2015).

The contributions of individual hitchhikers to MT plus end dynamics remain little understood (Akhmanova and Steinmetz, 2010; Akhmanova and Steinmetz, 2015; van de Willige et al., 2016). Reported mutant phenotypes tend to be subtle and often inconsistent, and their impact on MT dynamics may be indirectly influenced by additional cellular functions of these proteins (for example, APC acts as an actin nucleator) (Okada et al., 2010). In addition, significant differences were reported between +TIP behaviours in neuronal axons and nonneuronal cells, and careful validation is required before adapting the enormous body of knowledge acquired in non-neuronal cells to axonal contexts (Beaven et al., 2015).

XMAP215/CKAP5/CHTOG does not require EB proteins but can bind to MT plus ends via its N-terminal TOG domain (Fig. 2) (Al-Bassam and Chang, 2011). In this position, it acts as a processive microtubule polymerase by binding soluble tubulin heterodimers via its C-terminal TOG domains and promoting their polymerisation (Fig. 2) (Akhmanova and Steinmetz, 2015; Brouhard and Rice, 2014); this polymerisation activity can be regulated through mechanical stretch (Trushko et al., 2013). XMAP215 clearly plays a role in axons, for example during axon initiation and during growth cone guidance, and does so in close collaboration with the transforming acidic coiled-coil protein TACC3 (Bearce et al., 2015; Lowery et al., 2013; Nwagbara et al., 2014). Our own studies with depletion of Mini spindles (Msps), the *Drosophila* homologue of XMAP215, reveal a very severe reduction in axon growth (unpublished data); like in the case of EB1 depletion, this suggests a potential major role in MT polymerisation in axons.

As mentioned earlier, *in vitro* polymerisation assays have revealed a strong synergy between EB proteins and XMAP215 (Zanic et al., 2013). This seems surprising because observations *in vitro* and in HeLa cells suggest that both proteins are spatially separated (Fig. 2): XMAP215 binds preferentially to curved protofilaments occurring at the very tip, whereas the bulk of EB proteins binds up to several hundred nm away from the tip (Akhmanova and Steinmetz, 2015; Maurer et al., 2014). It has been proposed that EBs could contribute to the straightening of protofilaments at the MT plus end, thus helping XMAP215 to shift anteriorly to curved protofilaments and engage in its next polymerisation cycle (Brouhard and Rice, 2014; Zanic et al., 2013). Alternatively, it was reported in HeLa cells and cultured neurons that the protein Slain2 (with SxIP-like motifs) links EB1 to XMAP215 thus promoting MT polymerisation (van der Vaart et al., 2012; van der Vaart et al., 2011). A similar mechanism was suggested for the non-related Sentin protein in *Drosophila* which likewise contains SxIP-like motifs and links EB1 and Msps (Li et al., 2011; Li et al., 2012).

The MT plus end polymerisation machinery might contain additional factors redundant to XMAP215. For example, the EB-binding CLASP proteins contain TOG-like domains (Fig. 2); they regulate MT catastrophes and promote re-polymerisation via tubulin recruitment (Al-Bassam et al., 2010; Brouhard and Rice, 2014). In axons, CLASP function can switch between growth inhibition and promotion, crucially dependent on cellular contexts and GSK-3β signalling (Bearce et al., 2015). In a recent report, the reduction of axon growth observed upon CLASP knock-down in regenerating neurons was so strong (Hur et al., 2011) that CLASP might well act as a polymerase in this context. Furthermore, also stathmins might contribute as redundant factors during polymerisation because they bind tubulin heterodimers and can associate with MT plus ends (Fig. 2; see section 4).

Finally, it needs to be mentioned that some +TIPs primarily disassemble MTs (Homma et al., 2003). These include members of the kinesin-13 family (e.g. Kif2C/MCAK) which use their ATP-dependent motor domain to actively remove tubulins. They also include members of the kinesin-8 family which walk along MTs towards the plus tip and take tubulins with them when falling off the tip (Brouhard and Rice, 2014). Such mechanisms seem relevant in axons, as shown for the kinesin-13 member Kif2A which inhibits collateral branching (Homma et al., 2003).

## 6. Influence of lattice-based mechanisms on MT de/polymerisation

Axons are known to contain subpopulations of stabilised MTs (Baas et al., 2016). In our view the term stabilisation is imprecise and at least five modes of MT stabilisation need to be distinguished: (1) stabilising MTs through sustaining their plus end polymerisation (e.g. maintaining polymerase active; see previous section) (Qu et al., 2016); (2) stabilising MT lattices against depolymerisation (e.g. turning depolymerisation into pause or recovery); (3) stabilising MTs against disassembly through severing proteins (Yu et al., 2005); (4) stabilising MT minus ends through proteins such as CAMSAP/Patronin (Akhmanova and Hoogenraad, 2015); (5) stabilising MTs against mechanic deformation through bundling or other forms of cross-linkage.

In line with the topic of this review, we will focus here on a number of lattice-based mechanisms which affect plus end dynamics, either by preventing depolymerisation or through promoting polymerisation (see yellow arrow in Fig. 2). First, it has been reported that axonal MTs maintain high levels of GTP tubulin (Nakata et al., 2011) which, based on observations *in vitro*, should have stabilising functions. Thus, GTP tubulin at MT plus ends prevents catastrophes (section 3) and islands of GTP-bound tubulin within the MT lattice trigger rescue events (Dimitrov et al., 2008; Tropini et al., 2012).

Second, posttranslational modifications (PTM) of MTs are often taken as indicators for MT stability. In particular detyrosination and acetylation were shown to correlate well with stable MT fractions when analysed at the ultrastructural level (Baas et al., 2016). Commonly used indicators of acetylation are antibodies against acetylated K40 of α-tubulin. This residue is positioned on the luminal side of MTs (Howes et al., 2014; Li and Yang, 2015; Soppina et al., 2012); whether K40 itself mediates MT stabilisation, or other lysine residues potentially acetylated in parallel (Choudhary et al., 2009), will need further investigation. How PTMs stabilise MTs remains poorly understood: they could directly stabilise tubulin bonds, or they could modify MT interactions with stabilising MAPs or destabilising kinesins (Baas et al., 2016; Janke and Bulinski, 2011). Exploring these mechanisms is complicated by the pleiotropic functions of PTM-mediating enzymes; for example, the acetyltransferase α-TAT1 (responsible for K40 acetylation of α-tubulin) (Akella et al., 2010; Friedmann et al., 2012; Howes et al., 2014) regulates MT stability through direct MT interaction which is independent of its acetylation function (Kalebic et al., 2013).

Tubulin amination catalysed by transglutaminases, is a PTM essentially contributing to cold-stable MTs in brain extracts (Song et al., 2013). It has been suggested that amination could protect fragments of MTs from disassembly which could then be used as initiation points for *de novo* polymerisation, thus helping to maintain MT populations in axons (see Section 3) (Baas et al., 2016; Song et al., 2013). Apart from amination, cold protection is conferred also by MAP6/STOP (stable tubule only peptide; Fig. 2). MAP6 has been shown to bind and stabilise MTs preferentially at lower temperatures induced by conformational changes in its MT-binding Mc domain (Delphin et al., 2012; Poulain and Sobel, 2010). However, MAP6 seems to play roles also at normal temperatures, as suggested by progressive developmental brain dysfunction observed in MAP6 knock-out animals (Volle et al., 2013) (and references therein) - although these functions might work through mechanisms independent of MT binding (Deloulme et al., 2015).

Third, classical MAPs including MAP1B and tau have long been known as stabilisers against drug-induced de-polymerisation in axons (Takemura et al., 1992). They are likely to stabilise tubulin-tubulin bonds and/or compete for tubulins with destabilising kinesins or drugs. However, the mechanisms through which MAPs act remain little understood. For example, tau has been suggested to be co-assembled into MT lattices, whereas others suggest dynamic interaction with very brief dwell times on MTs (Amos and Schlieper, 2005; Janning et al., 2014; Kadavath et al., 2015); clearly, these two modes would be associated with very different mechanisms of MT stabilisation.

Further challenges for the understanding of classical MAPs are posed by the staggering inconsistency of phenotypes observed in different knock-out models of tau and MAP1B (Chilton and Gordon-Weeks, 2007; Morris et al., 2011; Villarroel-Campos and Gonzalez-Billault, 2014), as well as the enormous breadth of cellular functions they are suggested to perform, apart from stabilising MTs against depolymerisation. Such functions include MT bundling, protection against MT severing proteins, regulation of axonal transport, MT nucleation, actin regulation and MT-actin cross-linkage, the regulation of signalling processes and of MT plus end polymerisation (Elie et al., 2015; Morris et al., 2011; Rosenberg et al., 2008; Villarroel-Campos and Gonzalez-Billault, 2014).

Of these functions, potential roles in MT plus end polymerisation are of particular interest for this review, and several potential mechanisms have been proposed. First, MAP1B has been suggested to impact negatively on MT plus end polymerisation through sequestering EB1/3 in the cytoplasm (Tortosa et al., 2013). However, this report deviates from other reports showing that MAP1B promotes axon growth and MT polymerisation (Tymanskyj et al., 2012; Villarroel-Campos and Gonzalez-Billault, 2014). An alternative model can be deduced from work on the centrosomal protein TPX2 which localises along MT shafts up to their tips, where it directly binds and stabilises XMAP215 (Roostalu et al., 2015). Similarly, shaft-based tau was suggested to bind and stabilise EB1 (Sayas et al., 2015), and the same might be true also for MAP1B. Finally, *in vitro* work has shown that classical MAPs directly promote the incorporation of tubulins, potentially by neutralising or even attracting their negatively charged C-terminal domains (Lu and Luduena, 1994a) (and references therein).

A high degree of functional redundancy has been suggested for classical MAPs. Accordingly, the combined loss of different MAPs was shown to cause enhanced phenotypes including stronger reduction in neurite growth (Takei et al., 2000; Teng et al., 2001). More recently, proteins of the spectraplakin family, in particular ACF7/MACF1 and BPAG1/dystonin, were suggested as factors redundant to classical MAPs (Voelzmann et al., 2016). Like tau and MAP1B, spectraplakins are potent MT lattice binders and stabilisers, and they can also influence plus end polymerisation dynamics in axons (Alves-Silva et al., 2012; Qu et al., 2016; Sun et al., 2001). Unlike tau and MAP1B, spectraplakin deficiency causes very strong MT phenotypes correlating with neuro-developmental disorders as well as severe neurodegeneration in mouse and humans (Edvardson et al., 2012; Ferrier et al., 2013; Goryunov et al., 2010). Spectraplakins are therefore very likely candidates for masking key functions of tau and MAP1B and should be considered in future studies.

## 7. Other cellular functions of MT plus ends in axons

So far the focus of this review lay on the role of plus end de/polymerisation in the context of net growth and MT volume regulation. However, MT plus end dynamics have a greater scope of functions in axons. Analogous to the role that growth cones play for axons, plus ends can be considered as the steering units of MTs which can interact with their environment and determine the directionality and fate of MTs, thus turning MTs into important tools contributing to cellular processes. Below, we will depict some examples:

MT plus ends interact with proteins at the plasma membrane. Non-favourable membrane compartments contain factors repelling MTs by inducing their collapse. Such mechanism could be important in growth cones where MTs pointing away from the direction of future growth tend to retract (Bearce et al., 2015; Prokop et al., 2013). Some suggested factors mediating MT collapse include cortical katanin (Zhang et al., 2011) as well as Efa6 (the *C. elegans* homologue of PSD/pleckstrin and sec7 domains-containing protein) which is an inhibitor of axon regeneration (Chen et al., 2015; Chen et al., 2011; O’Rourke et al., 2010).

*Vice versa*, molecular networks associated with the cellular cortex can positively guide MT extension in neurons, for example by establishing links of plus ends to sub-membranous actin networks via spectraplakins (Alves-Silva et al., 2012) or drebrin-EB3 complexes (Geraldo et al., 2008). Furthermore, the MT plus end binding protein navigator1 localises to the periphery of growth cones and mediates axonal guidance (Martinez-Lopez et al., 2005).

MTs can also be captured at specialised membrane compartments, for example to deliver signals or materials via MT-based transport to cell junctions. Capture mechanisms were proposed to include the MT plus end binding proteins CLIP, APC, CLASP (Komarova et al., 2002; Schmidt et al., 2012; Swiech et al., 2011; Temburni et al., 2004), links from dynein to the NCAM adhesion receptor (Perlson et al., 2013), direct interaction of β3-tubulin with Netrin receptors (Qu et al., 2013), or palmitoylation of α-tubulin (Caron et al., 2001). As a further mode of mutual interaction, it has recently been described that cortical actin networks of axons act as powerful promoters of MT plus end polymerisation (Qu et al., 2016).

MTs are essential for the structure of endoplasmic reticulum (ER) which forms uninterrupted tubular networks all along axons that have been referred to as “neuron-within-neuron” (Berridge, 1998). The importance of axonal ER is illustrated by its close links to motor neuron disease (Blackstone et al., 2011; Yalçýn and O’Kane, 2013) and is explained by its many functions: ER regulates levels of intracellular free calcium in the axonal cytoplasm, calcium homeostasis, fission and fusion of mitochondria, and it has been suggested to mediate local translation of membrane-associated or releasable proteins as well as lipid biogenesis contributing to axonal membranes (Berridge, 1998; Friedman and Nunnari, 2014; Gonzalez and Couve, 2014; Korobova et al., 2013; Wojnacki Fonseca and Galli, 2016). The SxIP protein STIM1 can link ER to MT plus ends to mediate directed ER elongation (Grigoriev et al., 2008). Since STIM1 can also regulate calcium influx at the plasma membrane (Cahalan, 2009), it might be a crucial mediator between ER and axonal surfaces to control levels of intracellular free calcium, and MTs might play important roles during this process. This latter notion would be in line with the general idea that MTs, and in particular also MT plus ends, are important hubs for signalling events in cells (Dent and Baas, 2013).

## 8. Conclusions and future prospects

Fig. 2 illustrates the complexity of the various regulatory networks coordinating MT plus end dynamics in axons. To make sense of these dynamics at the cellular level, we need to integrate the detailed knowledge we have gathered, into conceptual systemic understanding. For example, as was explained above, a number of factors including EB1, stathmins, XMAP215, CLASPs, TBCs and tubulins themselves have the ability either to promote or to inhibit MT plus end polymerisation. This cocktail of bi-directional regulators could form an effective buffer against unwanted changes in polymerisation processes. On the other hand, intentional changes induced by signals need to achieve a systemic reversal of the activity status of many or even all of these regulators. Furthermore, a number of proteins mentioned in this review are not restricted to one function, but can act through very different mechanisms. Extreme examples are the classical MAPs (section 5) and the multi-functional cylindromatosis protein (CYLD), which directly interacts with MTs, binds to MT plus ends through EB proteins, regulates MT deacetylation and acts as a de-ubiquitinase (Yang and Zhou, 2016). Therefore, a reductionist approach by studying single factors in isolation seems insufficient to explain MT behaviours in cellular environments.

A systemic view of the problem of MT dynamics can be of benefit also by facilitating the discovery of commonalities and fundamental principles. For example, the different stages of a neuron’s life, as depicted in Fig. 1, illustrate the continuum of the same MT bundle from the birth of the axon through to its degeneration. As already explained in section 2, MT de/polymerisation occurs throughout all these stages, and it seems reasonable to assume that the same fundamental machinery of MT de/polymerisation underlies processes of growth, branching, degeneration and regeneration. Accordingly, factors like tau, spectraplakins or CLASP to name but a few, play important roles during development as well as during the adult life of axons (Hur et al., 2011; Prokop et al., 2013). Mechanistic understanding gained from work on one stage or functional context might therefore apply more widely to studies of axonal biology.

Building on this idea, we started to assess whether MT regulating mechanisms known from axon development might play similar roles during axonal ageing. This work led us to propose the model of local homeostasis of axons (Fig. 3) (Prokop, 2016). In this model, we propose that MTs of mature axons are constantly renewed through steady state de/polymerisation processes (see also section 2). However, MTs do not necessarily align into bundles: the force environment in axons, which is rich in kinesins and dyneins, causes MTs to curl, and excessive looping of MTs is the most common phenotype we observe upon manipulations of different classes of MT regulators (Fig. 3) (Beaven et al., 2015; Sánchez-Soriano et al., 2009). These loops and their diameters are comparable to loops observed in gliding assays *in vitro* where MTs are kept on kinesin carpets (Liu et al., 2011). Models explaining this phenomenon have been proposed (Ziebert et al., 2015). The tendency of axonal MTs to go into disarray requires that different classes of MT regulators impose order by arranging axonal MTs into bundles and maintaining this condition for decades (details in Fig. 3). We predict that mutations of single regulators do not eliminate but weaken this machinery, increasing the probability of accidental MT disorganisation. This could explain axonal swellings observed in neurodegenerative disorders associated with various MT regulators (as well as in normal ageing). These swellings are thought to form traps for organelles and initiation points of axon degeneration (Adalbert and Coleman, 2012; Prokop et al., 2013).

**Figure 3:**
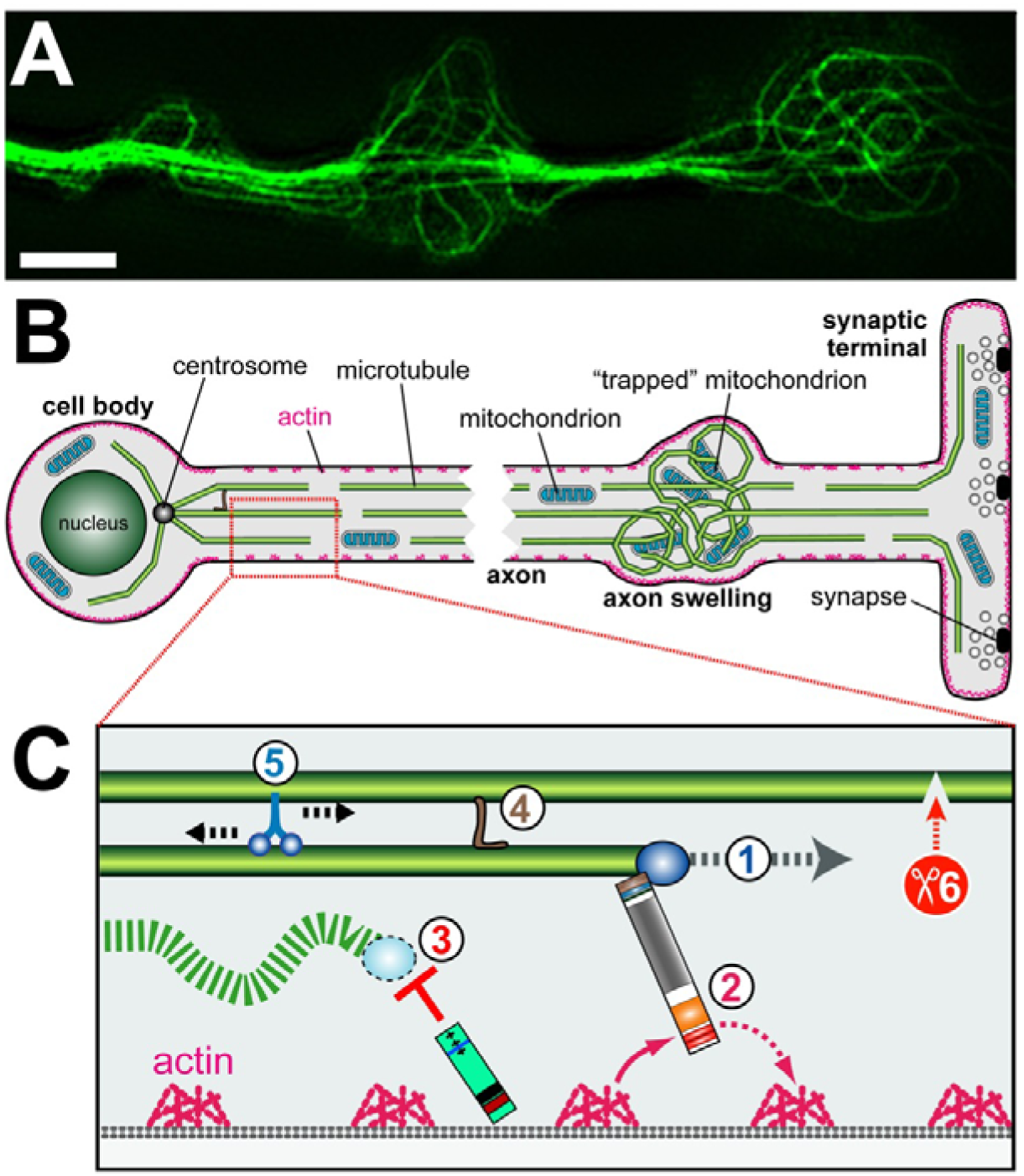
The model of local homeostasis in axons. **A**) MT looping in axons of primary neurons is a common phenotype observed upon mutation of MT regulators (scale bar 1µm). **B**) MT looping observed in primary neurons might reflect the same pathomechanisms that give rise to axon swellings during ageing or neurodegeneration. **C**) The model of local homeostasis proposes that continued polymerisation of MTs **(1)** poses a constant risk of MT disorganisation, and that different MT regulators contribute in different ways to maintain MT bundles and prevent axon swellings: spectraplakins **(2)** bind MT plus ends and guide them along cortical F-actin to lay them out into parallel bundles (Alves-Silva et al., 2012); cortical collapse factors **(3)** act as check points, capturing off-track MTs and eliminating them (unpublished results); bundling mechanisms **(4)** are likely to stabilise MT bundles from within; MT sliding **(5)** could ensure even MT densities along the axon shaft; **(6)** MT severing could provide an important mechanisms to ensure MT turn-over in order to prevent senescence.

This model represents an attempt to explain axon biology through integrating our knowledge about regulatory mechanisms of MTs. It provides a hypothesis that can be experimentally tested and refined. Such experiments require cellular systems in which the various MT regulators and their interactors can be functionally manipulated in parallel, in order to assess functional redundancies and study the interfaces between different regulatory mechanisms. As has been detailed elsewhere (Prokop et al., 2013), neurons of the fruitfly *Drosophila* provide the means to carry out such studies. *Drosophila* has been used for decades to unravel biological complexity because it provides access to a powerful combination of highly efficient genetic and experimental approaches (Hahn et al., 2016). Importantly, many of the lessons learned in the fly can be translated into mammalian systems, and this clearly applies to cytoskeletal studies in axons (Beaven et al., 2015; Prokop et al., 2013; Sánchez-Soriano et al., 2009).

## 9. Acknowledgement

AV is supported by a postdoctoral fellowship of the German Research council (DFG; VO 2071/1-1), NS by a BBSRC grant (BB/M007456/1), SPP by an Early Career Fellowship from the Leverhulme Trust (ECF-2015-229), IH and AP by a BBSRC grant (BB/L000717/1), and AP by a BBSRC grant (BB/M007553/1).

